# An anti-ACVR1 antibody exacerbates heterotopic ossification by fibro/adipogenic progenitors in fibrodysplasia ossificans progressiva mice

**DOI:** 10.1101/2021.07.23.451471

**Authors:** John B. Lees-Shepard, Sean J. Stoessel, Julian Chandler, Keith Bouchard, Patricia Bento, Lorraine N. Apuzzo, Parvathi M. Devarakonda, Jeffrey W. Hunter, David J. Goldhamer

## Abstract

Fibrodysplasia ossificans progressiva (FOP) is a rare genetic disease characterized by progressive and catastrophic heterotopic ossification (HO) of skeletal muscle and associated soft tissues. FOP is caused by dominantly acting mutations in the bone morphogenetic protein (BMP) type I receptor, ACVR1 (also known as ALK2), the most prevalent of which is an arginine to histidine substitution [ACVR1(R206H)] in the glycine-serine rich intracellular domain of the receptor. A fundamental pathological consequence of FOP-causing ACVR1 receptor mutations is to enable activin A to initiate canonical BMP signaling in responsive progenitors, which drives skeletogenic commitment and HO. With the clear targets of activin A and ACVR1 identified, development of antibody therapeutics to prevent ligand-receptor interactions is an interventional approach currently being explored. Here, we developed a monoclonal blocking antibody (JAB0505) to the extracellular domain of ACVR1 and tested its ability to inhibit HO in established FOP mouse models. JAB0505 inhibited BMP-dependent gene expression in wild-type and ACVR1(R206H)-overexpressing cell lines. Strikingly, however, JAB0505 treatment markedly exacerbated injury-induced HO in two independent FOP mouse models in which ACVR1(R206H) was either broadly expressed, or more selectively expressed in fibro/adipogenic progenitors (FAPs). JAB0505 drove HO even under conditions of activin A inhibition, indicating that JAB0505 has receptor agonist activity. JAB0505-treated mice exhibited multiple, distinct foci of heterotopic lesions, suggesting an atypically broad anatomical domain of FAP recruitment to endochondral ossification. In addition, skeletogenic differentiation was both delayed and prolonged, and this was accompanied by dysregulation of FAP population growth. Collectively, alterations in the growth and differentiative properties of FAPs and FAP-derived skeletal cells are implicated in the aggravated HO phenotype. These data raise serious safety and efficacy concerns for the use of anti-ACVR1 antibodies to treat FOP patients.

## Introduction

Fibrodysplasia ossificans progressiva (FOP) is a rare, autosomal-dominant, genetic disease of progressive heterotopic ossification (HO) that primarily affects skeletal muscle and associated connective tissues. The cumulative effect of HO, which typically begins in early childhood (1), leads to progressive immobility and shortened lifespan (2). The most common FOP mutation in *Acvr1* results in an arginine to histidine amino acid substitution at position 206 in the intracellular glycine-serine domain of ACVR1 [ACVR1(R206H)] (3), which renders this bone morphogenetic protein (BMP) receptor responsive to activin ligands (4–7) and hypersensitive to select BMP ligands (8–11). While the physiological relevance of hypersensitivity to BMP ligands is unclear, blocking antibodies to activin A are highly effective at inhibiting HO in preclinical mouse models of FOP, and a clinical trial is underway to evaluate this strategy (NCT03188666). Here, we address whether an antibody directed against the extracellular domain of ACVR1(R206H) that interferes with ligand-dependent receptor activation would inhibit HO and constitute a potential therapeutic approach for FOP. We present the unexpected finding that an anti-ACVR1 antibody (JAB0505) that effectively blocks ligand-dependent osteogenic reporter gene expression in cultured C2C12 cells profoundly exacerbates HO in FOP mouse models. As our previous work and that of others indicates that fibro/adipogenic progenitors (FAPs) are a key causative cell type in FOP (6, 12, 13), we addressed how antibody treatment affects BMP signaling, expansion, and differentiation of ACVR1(R206H)-expressing FAPs (R206H-FAPs). We demonstrate that JAB0505 is a weak agonist of ACVR1(R206H) and can activate osteogenic signaling in the absence of activin A. Treatment of FOP mice with JAB0505 altered the expansion, recruitment, and osteogenic differentiation kinetics of R206H-FAPs, all of which likely conspire to dramatically enhance HO. Recently, independent studies have confirmed extreme worsening of HO with anti-ACVR1 antibodies (14). Together, these data indicate that blockade of ligand-receptor interactions by the use of bivalent anti-ACVR1 monoclonal antibodies is not a viable therapeutic approach for FOP.

## Results and Discussion

### Characterization of the anti-ACVR1 monoclonal antibody, JAB0505

A mouse anti-ACVR1 monoclonal antibody (mAb) was isolated from an immune-biased phage-display antibody library that was constructed from mice immunized with DNA encoding the extracellular domain of ACVR1. This antibody was affinity matured by PCR-mutagenesis of the complementary determining regions (CDRs) to yield antibody JAB0505. Affinity maturation improved the ability of JAB0505 to inhibit ligand-induced ACVR1 signaling *in vitro* (Supplemental Fig. 1). The equilibrium dissociation constant (K_D_) of JAB0505 for binding to ACVR1 was determined to be ∼2.5 X 10^-8^ M by surface plasmon resonance (Supplemental Fig. 1a). Inhibition of BMP-dependent signaling by JAB0505 was demonstrated *in vitro* using mouse C2C12 cells that endogenously express wild-type (WT) ACVR1. Quantification of BMP type I receptor mRNA abundance in C2C12 cells by RT-qPCR revealed slightly higher levels of ALK3 (BMPR1A) than ACVR1 (ALK2), much lower levels of ALK1, and no detectable ALK6 mRNA (Fig. 1a). Cell surface expression of ACVR1 and ALK3 on C2C12 cells was confirmed by flow cytometry with receptor-specific antibodies (Fig. 1b). JAB0505 does not recognize ALK3, as evidenced by the loss of cell surface binding of JAB0505 to ACVR1-KO C2C12 cells (Fig. 1c). The ability of JAB0505 to block ligand-dependent BMP signaling was assessed in wild-type and ACVR1(R206H)-transfected C2C12 cells by quantifying activity of the Id1-luciferase reporter, BRE-luc, a well-established transcriptional readout of BMP signaling activity (15). JAB0505 completely blocked BMP6-, BMP7-, BMP9-, and BMP10-induced luciferase activity in the wild-type C2C12-BRE-Luc reporter cell line (Fig. 1d; Supplemental Fig. 1d-f). In the ACVR1(R206H)-transfected C2C12-BRE-Luc reporter cell line, JAB0505 did not diminish luciferase activity in response to BMP2 or BMP4 (Supplemental Fig. 1g, h), which signal principally through ALK3 (16), but blocked hyper-responsive signaling to BMP9 (Fig. 1d; Supplemental Fig. 1b). Although BMP9 also signals via ALK1, the return to baseline with JAB0505 treatment, and the very low expression level of ALK1 in this cell line (Fig. 1a), strongly suggests that the observed BMP9-dependent signaling is almost exclusively mediated by ACVR1. Baseline luciferase activity was substantially higher than that observed for wild-type reporter cells and unaffected by JAB0505 (Fig. 1d), probably reflecting either weak, ligand-independent signaling due to overexpression of the mutant receptor, or differential responsiveness to ligands in the growth serum.

**Figure 1.**
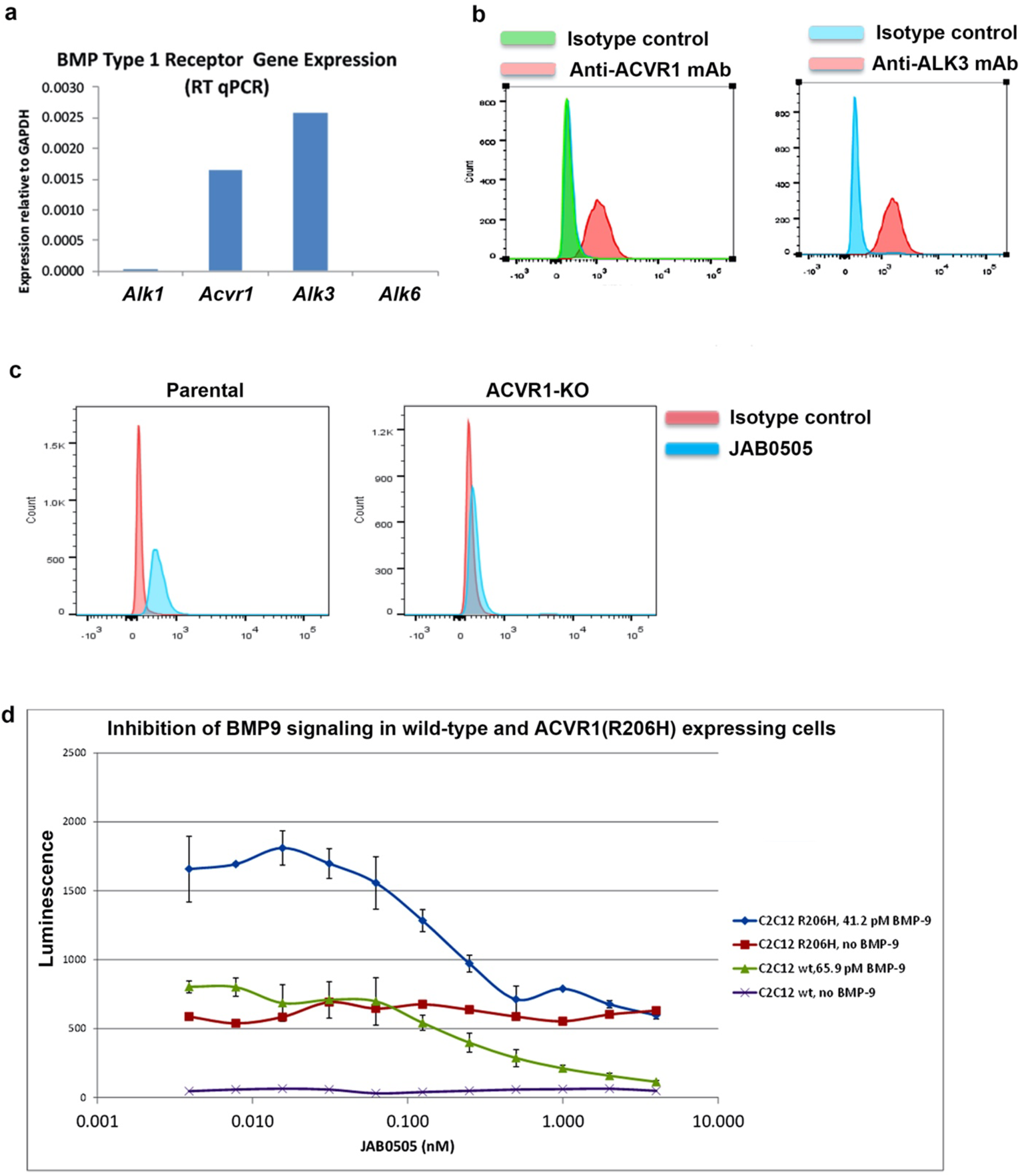
JAB0505 is an ACVR1-blocking monoclonal antibody. (**a**) Mouse myoblast C2C12 cells express *Acvr1* and *Alk3* but not *Alk1* or *Alk6*, as detected by RT-qPCR. **(b)** Surface expression of ACVR1 and ALK3 on C2C12 cells, as detected by flow cytometry. **(c)** Monoclonal antibody JAB0505 binds to parental C2C12 cells, but not *Acvr1* KO cells, as assessed by flow cytometry. **(d)** JAB0505 inhibits BMP signal activation in wild-type and ACVR1(R206H)-overexpressing C2C12 cells in a dose-dependent manner, as determined by quantification of BRE-luciferase activity.

### JAB0505 dramatically exacerbates and prolongs HO in FOP mice

We next tested the ability of JAB0505 to inhibit injury-induced HO in FOP mouse models. First, we used conditional *Acvr1^FLEx(R206H)/+^*; CAG-Cre^ERT2^ mice in which recombination of the *Acvr1^R206H^* allele is driven by the ubiquitously expressed CAG-Cre driver and is tamoxifen-dependent. Following tamoxifen administration and a 5-to-7-day washout period, HO was induced by intramuscular injection of cardiotoxin into the gastrocnemius muscle. Surprisingly, administration of 10 mg/kg JAB0505 at the time of muscle injury dramatically exacerbated the formation of heterotopic bone, as assayed at day 14 (data not shown) and day 20 (Fig. 2a; n=4) post injury. To further investigate the mechanism of exacerbated heterotopic bone formation, we tested the effects of JAB0505 in *Acvr1^tnR206H^* mice (6), in which the Tie2-Cre driver (17) was used to target *Acvr1^R206H^* expression to FAPs, a major cell-of-origin in HO (6, 12, 18). In agreement with the observations in globally recombined *Acvr1^FLEx(R206H)/+^*;CAG-Cre^ERT2^ mice, a single intraperitoneal injection of 10 mg/kg JAB0505 at the time of muscle pinch injury profoundly exacerbated HO in *Acvr1^tnR206H/+^*;Tie2-Cre FOP mice, as revealed by μCT (Fig. 2b). Neither uninjured FOP mice nor control mice lacking either Tie2-Cre or the *Acvr1^tnR206H^* allele developed HO in response to JAB0505 (data not shown).

**Figure 2.**
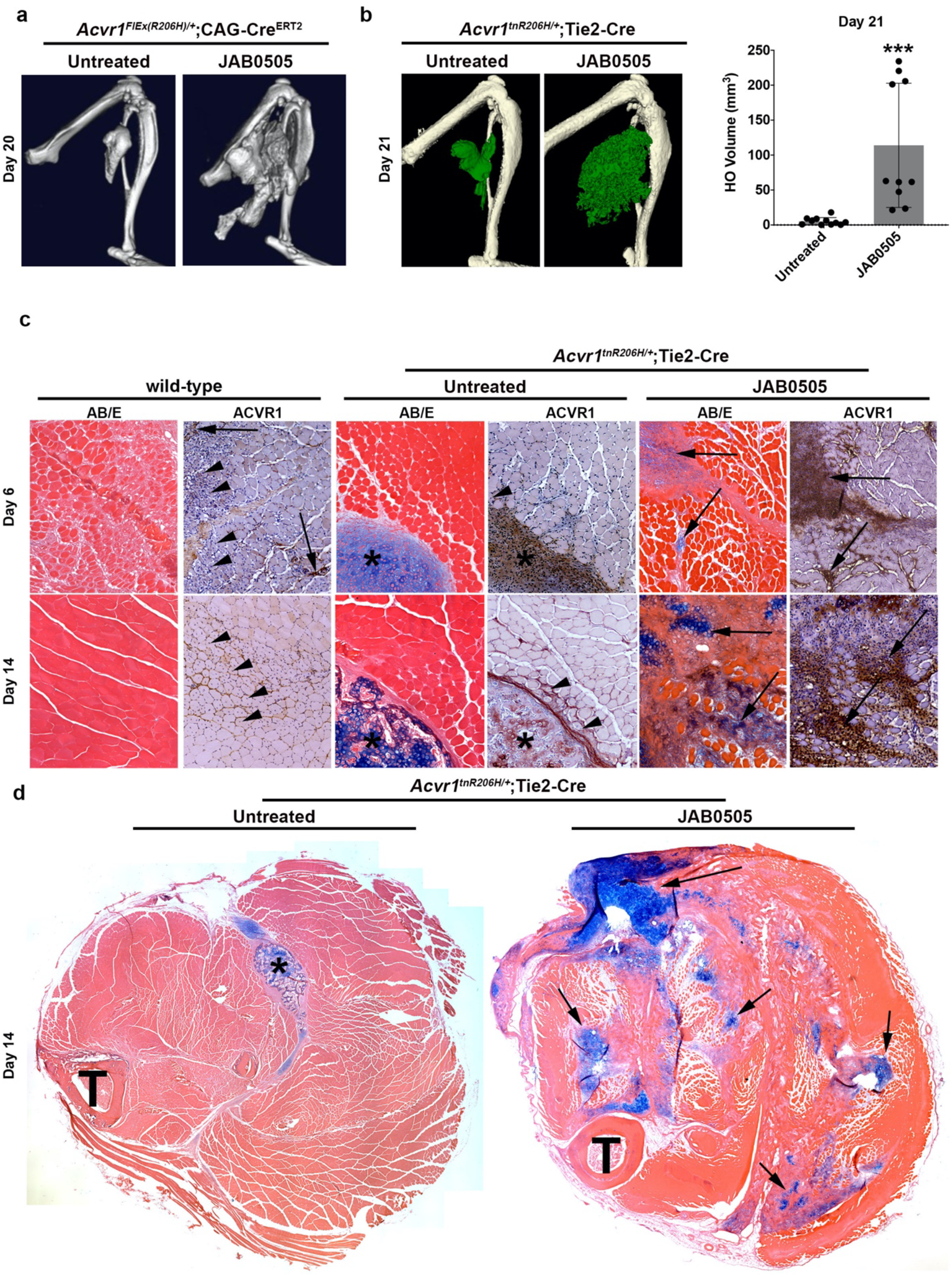
JAB0505 profoundly exacerbates HO in FOP mice. (**a**) Representative µCT images of HO in *Acvr1^FLEx(R206H)/+^*;CAG-Cre^ERT2^ mice 20 days after cardiotoxin-induced injury of the gastrocnemius muscle (Untreated, n = 3; JAB0505, n = 4). **(b)** Representative µCT images and volumetric quantification of HO (pseudocolored green) in *Acvr1^tnR206H/+^*;Tie2-Cre mice 21 days after pinch injury of the gastrocnemius muscle (Untreated, n = 11; JAB0505, n = 10. Bars represent ± SD. p = 0.001, unpaired t-test). **(c)** Alcian blue staining for cartilage (arrows) and immunohistochemical staining of ACVR1 (brown, arrows) on cross-sections of pinch-injured muscle from untreated wild-type mice, untreated *Acvr1^tnR206H/+^*;Tie2-Cre mice (lesional region denoted by asterix), and JAB0505-treated *Acvr1^tnR206H/+^*;Tie2-Cre mice. Centrally located myofiber nuclei (arrowheads) identify regenerated fibers, which were abundant in wild-type mice, rare in *Acvr1^tnR206H/+^*;Tie2-Cre mice, and undetected in JAB0505-treated *Acvr1^tnR206H/+^*;Tie2-Cre mice. AB/E, Alcian blue/eosin. Sections processed for immunohistochemistry were counterstained with hematoxylin. **(d)** Low magnification images of lower hindlimb cross-sections of *Acvr1^tnR206H/+^*;Tie2-Cre mice at day 14 post-injury. As compared to HO (asterix) of untreated *Acvr1^tnR206H/+^*;Tie2-Cre mice at this stage, boney HO was less prevalent in JAB0505-treated *Acvr1^tnR206H/+^*;Tie2-Cre mice, which also exhibited broader distribution of cartilaginous HO (arrows). Sections were stained with Alcian blue and eosin.

To assess the kinetics of bone formation in JAB0505-treated *Acvr1^tnR206H/+^*;Tie2-Cre FOP mice, μCT imaging was performed at weekly intervals, from 2 to 5 weeks post-injury. In untreated *Acvr1^tnR206H/+^*;Tie2-Cre mice, μCT imaging showed that mineralized, electron-dense, heterotopic bone commonly reaches its maximum extent and density by approximately day 14 (Fig. 2a). Notably, the time course of mature, mineralized heterotopic bone formation is substantially prolonged in these JAB0505-treated FOP mice. This is evident at day 14 post-injury, when most of the radiographically detectable tissue is below the threshold set for mineralized bone (Supplemental Fig. 2). Indeed, the overt heterotopic bone is often scattered or lacy in appearance by μCT imaging and comprises only a small fraction of the lesional area at this time (Supplemental Fig. 2). By 21 days post-injury, mineralized lesions had grown substantially in JAB0505-treated mice, with an average bone volume approximately 20-fold greater than untreated FOP mice (Fig. 2b). Remarkably, bony lesions of JAB0505-treated mice continued to grow up through the experimental endpoint of 35 days post-injury (Supplemental Fig. 2), although quantification was impractical due to the intimate association of heterotopic bone with the limb skeleton. These data suggest that the mechanisms that normally render heterotopic bone growth self-limiting are inoperative or severely impaired in JAB0505-treated mice.

### A broadened domain of cell recruitment and delayed skeletal differentiation contribute to the pathology of JAB0505-treated Acvr1^tnR206H/+^*;Tie2-Cre* FOP mice

Histological analyses revealed additional differences in response to muscle injury between JAB0505-treated and untreated *Acvr1^tnR206H/+^*;Tie2-Cre FOP mice. Muscle pinch injury of untreated FOP mice results in the development of histologically identifiable cartilage by days 5-6, and the subsequent gradual replacement of cartilage with bone beginning at approximately day 10 (Fig. 2c; unpublished observations). Following injury, lesional growth typically proceeds from a primary area of mesenchyme accumulation, ultimately leading to heterotopic cartilage and bone that is clearly demarcated from surrounding muscle tissue not actively engaged in skeletogenesis (6). By day 6 post-injury, cartilage lesions of *Acvr1^tnR206H/+^*;Tie2-Cre mice are well-defined and stain intensely with Alcian blue, which binds acidic mucopolysaccharides and glycoproteins of the cartilage matrix (Fig. 2c). This contrasts with JAB0505-treated mice, in which cartilage lesions at day 6 had a less mature morphology and stained weakly with Alcian blue (Fig. 2c), indicating that the formation or maturation of cartilage is delayed by JAB0505 treatment. Also apparent was a delay in the appearance of mineralized bone at day 14 post-injury. At this stage, lesions of JAB0505-treated mice predominantly consisted of cartilage, in contradistinction to untreated FOP mice in which lesions at this stage were primarily comprised of mineralized bone, sometimes with residual hypertrophic cartilage remaining (Fig. 2c, d)(6). These histological observations indicate that slower kinetics of cartilage differentiation and transition to bone are, at least partially, responsible for prolonging the time course of HO. Strikingly, whereas day 14 lesions of untreated FOP mice were typically comprised of a contiguous bony mass, numerous Alcian blue-positive cartilage foci were scattered throughout much of the hindlimb musculature of JAB0505-treated mice after a single pinch of the gastrocnemius muscle (Fig. 2c, d). Probably due to the more widely disseminated pathological response of JAB0505-treated mice, muscle integrity was severely diminished, with only scattered muscle fibers remaining in some areas (Fig. 2c, d).

The difference in the anatomical distribution of cartilage at day 14 was reflected in the pattern of ACVR1 protein expression at day 6, when small foci of high ACVR1 expression were widely distributed in injured muscle of JAB0505-treated mice (Fig. 2b). We previously showed that ACVR1 protein expression in FOP mice is upregulated shortly after injury in cells of interstitial regions that co-express the chondrogenic marker, SOX9, and marks heterotopic skeletal tissues at later stages (6). Based on these observations, we have proposed that these connective tissue regions are areas of skeletal progenitor cell recruitment that help drive lesional growth (6). Collectively, these data suggest a model whereby treatment with JAB0505 exacerbates HO in FOP mice by expanding the anatomical domain and temporal window of FAP recruitment to skeletogenic lineages, and consequently, of FAP-driven HO.

Despite its potency in exacerbating HO, JAB0505 alone was not sufficient to trigger heterotopic bone formation in the absence of injury. Given the more widely disseminated disease that characterizes JAB0505-treated FOP mice, it is reasonable to suggest that JAB0505 reduces the injury threshold required to drive R206H-FAPs (and perhaps other progenitors) to HO, thereby resulting in an anatomically broader and sustained engagement of skeletal progenitors in heterotopic lesion formation and growth. To address this possibility, we tested whether JAB0505 can amplify a minor, sub-clinical injury that alone does not cause HO. To this end, the effects of JAB0505 were assessed after the tibialis anterior muscle of *Acvr1^tnR206H/+^*;Tie2-Cre mice was injected with 50 μL of 2.5% methylcellulose, which elicits no HO in this FOP mouse model (Supplemental Fig. 3) (6). Remarkably, administration of JAB0505 on the day of methylcellulose injection resulted in a dramatic HO response that was similar in magnitude to its effect following muscle pinch or cardiotoxin injection (Supplemental Fig. 3). As tissue inflammation is a trigger for HO in FOP (19, 20), one consequence of JAB0505 treatment may be to amplify and sustain the inflammatory response resulting from muscle injury, thereby amplifying what is otherwise an insufficient injury to induce R206H-FAPs to engage in endochondral bone formation in methylcellulose-treated limbs, or when located at a distance from a localized pinch injury. Studies by Rando and colleagues (21) predict that R206H-FAPs exposed to a mild injury would leave quiescence and enter a transitional and normally adaptive cell state called G_alert_, which positions stem cells to respond rapidly to injury and stress. This cell state, which is induced by systemic injury factors (21), presumably would be insufficient for R206H-FAPs to respond to JAB0505, as HO is restricted to the injured limb whereas both R206H-FAPs and JAB0505 are broadly distributed. In this context, an amplified inflammatory response caused directly or indirectly by JAB0505 could provide the additional stimulus for limb-specific and robust HO driven by R206H-FAPs. Alternatively, injection of a viscous carrier such as methycellulose may provide a stimulus that is sufficient to move R206H-FAPs of the injured limb to an activated state beyond G_alert_ in which cells are poised to enter the endochondral pathway as a direct response to JAB0505 activating ACVR1(R206H) complexes on FAPs, independent of JAB0505’s effect on tissue inflammation. Evidence that JAB0505 both potentiates the inflammatory response after injury and acts as a receptor agonist are presented below.

### The effects of JAB0505 on skeletal differentiation and ACVR1 activation are ligand, cell, and context-dependent

We next explored the relative responsiveness of wild-type and R206H-FAPs to activin A and BMP6, singly and in combination with JAB0505. JAB0505 was effective at blocking BMP6-induced chondrogenic and osteogenic differentiation of wild-type FAPs (Supplemental Fig. 4a), but only partially reduced pSMAD1/5/8 levels in BMP6-treated cells, despite molar ratios of JAB0505:ligand of 70:1 and higher (Supplemental Fig. 4a, b). Although, JAB0505 reduced activin A-driven SMAD1/5/8 phosphorylation of R206H-FAPs to a comparable degree (Supplemental Fig. 4b), it did not appreciably inhibit skeletal differentiation (Fig. 3a, b). The apparent disparity between effects of these ligands, and between short-term assays of receptor activation and long-term differentiation assays, suggests the engagement of multiple receptors, pathways, or both, in a context-dependent manner. In fact, BMP6 interacts with the SMAD1/5/8-activating receptors ALK3 and ALK6 (22), both of which are expressed by FAPs (unpublished observations). Further, knockout of ACVR1 (ALK2) in mouse embryonic fibroblasts inhibited chondrogenic differentiation in micromass cultures, even though pSMAD1/5 was induced, presumably through ALK3 (10). That efficient blockade of ligand-induced ACVR1 receptor activation by JAB0505 in ACVR1(R206H)-expressing C2C12 cells (Fig. 1d) does not predict the effects of JAB0505 in FAPs, highlights the complexity of BMP signaling and demonstrates the importance of validating such data in cells relevant to FOP.

**Figure 3.**
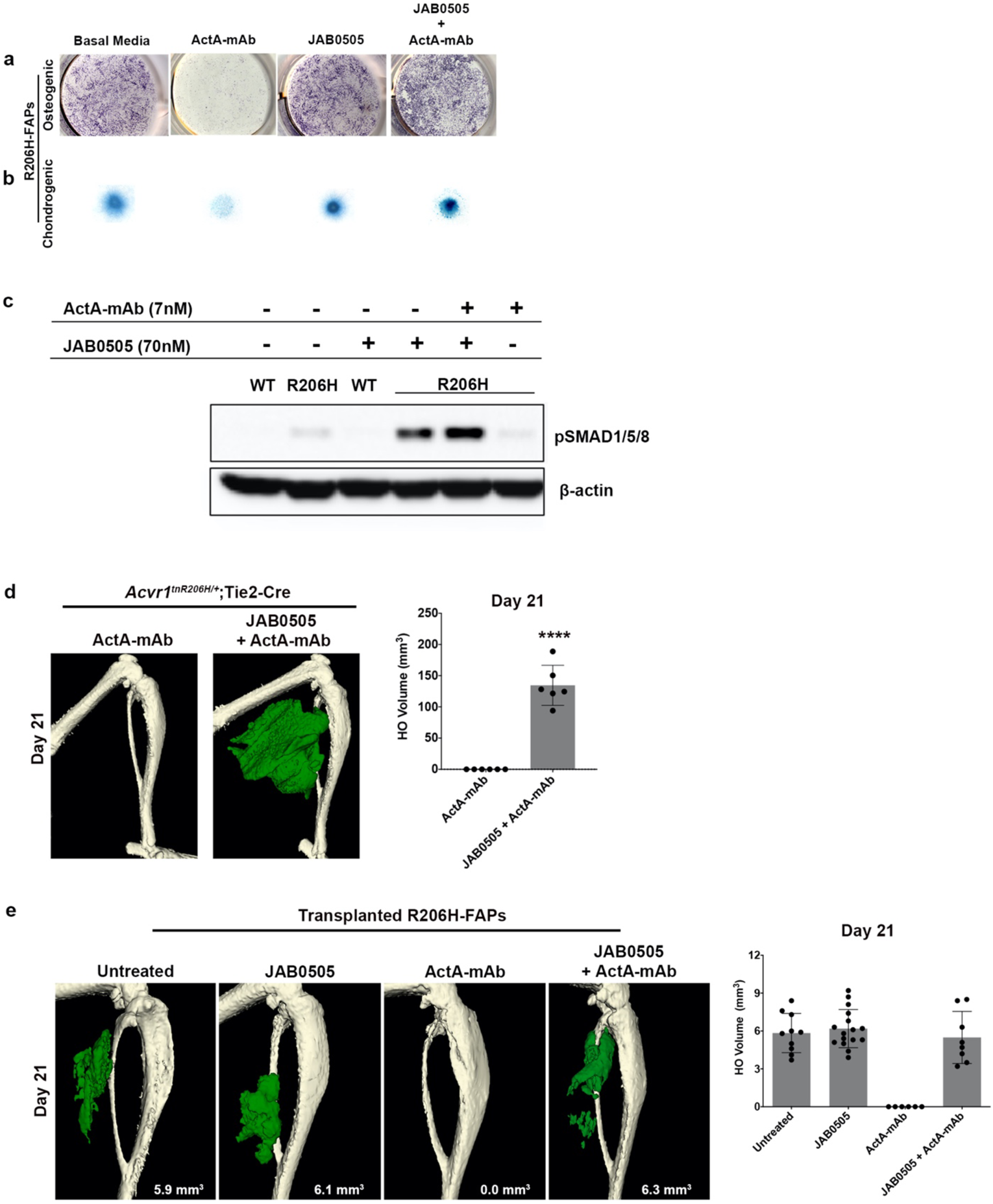
JAB0505 agonizes signaling by ACVR1(R206H). (**a**) Osteogenic differentiation of monolayer R206H-FAP cultures, as assessed by ALP staining (purple), and **(b)** chondrogenic differentiation of micromass culture assessed by Alcian blue staining. ActA-mAb was used at 1 μg/mL (7 nM) and JAB0505 was used at 10 μg/mL (∼70 nM). **(c)** Western blot of phosphorylated SMADs1/5/8 (pSMAD1/5/8) in wild-type (WT) and R206H-FAPs (R206H). β-actin was used as a loading control. **(d)** µCT of the distal hindlimb of *Acvr1^tnR206H/+^*;Tie2-Cre mice at day 21 post-injury. At the time of muscle injury, mice were treated with ActA-mAb (10 mg/kg) alone or ActA-mAb with JAB0505 (10 mg/kg). HO is pseudocolored green, and quantification is shown. ActA-mAb, n = 6; JAB0505 + ActA-mAb, n = 6. Bars represent ± SD; p < 0.0001, unpaired t-test. **(e)** µCT of the distal hindlimb at day 21 post-transplantation of R206H-FAPs into the injured gastrocnemius of SCID hosts. ActA-mAb (10 mg/kg) and JAB0505 (10 mg/kg) were administered at the time of transplantation. HO is pseudocolored green and quantified, with bars representing ± SD. Untreated, n = 10; JAB0505, n = 16; ActA-mAb, n = 6; JAB0505 + ActA-mAb, n = 8.

### JAB0505 functions as a weak agonist of ACVR1(R206H) in FAPs

As activin A is an obligatory ligand for HO formation in FOP mice (4, 6), we sought to test whether the effects of JAB0505 are dependent on activin A in culture and *in vivo*. As previously shown (6, 12), R206H-FAPs, but not wild-type FAPs, undergo a low level of osteogenic and chondrogenic differentiation without addition of exogenous ligand, a response that is completely blocked by anti-activin A antibody, indicating that activin A is present in the culture media (Fig. 3a, b). Notably, JAB0505 is sufficient to drive a comparable degree of chondrogenic and osteogenic differentiation of R206H-FAPs when serum activin A is neutralized with an anti-activin A monoclonal antibody (ActA-mAb) (Fig. 3a, b), demonstrating that JAB0505 functions as a weak receptor agonist in this setting. Western blot analysis of SMAD1/5/8 phosphorylation confirmed activation of BMP signaling in R206H-FAPs by JAB0505 in the absence of activin A (Fig. 3c), and this was also observed when JAB0505 was used at the same concentration as activin A (1 nM; data not shown). JAB0505 did not induce skeletal differentiation or SMAD1/5/8 phosphorylation of wild-type FAPs (Fig. 3c; Supplemental Fig. 4a), indicating that its agonist activity is functionally similar to activin A in that it activates BMP signaling only through ACVR1(R206H). This weak agonist activity can also partially explain the inability of JAB0505 to block activin A-induced skeletogenic differentiation and SMAD1/5/8 phosphorylation in cultured R206H-FAPs (Supplemental Fig. 4a, b).

We next tested whether JAB0505 can function as a receptor agonist in FOP mice. The gastrocnemius muscle of *Acvr1^tnR206H/+^*;Tie2-Cre FOP mice was pinch injured, and mice were treated with 10 mg/kg JAB0505 at the time of injury, with or without 10 mg/kg ActA-mAb, a dose that blocks injury-induced HO with 100% efficacy in this model (6). FOP mice treated with both JAB0505 and ActA-mAb also developed explosive HO (Fig. 3d), indicating that JAB0505 can replace the essential functions of activin A in driving the formation and growth of injury-triggered HO. Further, the results highlight an apparent paradox, namely, that receptor activating activities that are demonstrably weaker than those of activin A in cell culture assays (Fig. 3a-c), nevertheless profoundly worsen HO relative to the activin A-driven process in untreated FOP mice. It is reasonable to suggest that the slower kinetics and prolonged period of HO progression in these JAB0505-treated mice, which is correlated with exacerbation of HO, is at least partially due to the ability of JAB0505 to weakly activate ACVR1(R206H) while blocking productive interactions between ACVR1 and activin A, which more efficiently drives skeletal differentiation. However, signal potency is unlikely to be the sole factor dictating HO severity. Instead, temporal and spatial distribution of ACVR1 signal activation within injured muscle may also regulate HO severity. Indeed, the observations of broader engagement of muscle resident skeletal progenitors and a prolonged period of HO progression are consistent with broad spatial distribution of a weakly activating but long-acting agonist antibody, such as JAB0505.

As cells in addition to FAPs are recombined in *Acvr1^tnR206H/+^*;Tie2-Cre mice, including endothelial cells and CD45+ hematopoietic cells (6, 17, 18, 23), we further assessed FAPs as a target of receptor agonist properties of JAB0505 in transplantation assays, FAPs were isolated, expanded in culture, and transplanted into the pre-injured gastrocnemius muscle of SCID mice, as previously described (6, 12). Transplanted R206H-FAPs consistently formed heterotopic bone, as assayed by μCT at day 21 post-transplantation (Fig. 3e; n=10), and injection of ActA-mAb on the day of transplantation completely blocked osteogenic differentiation (Fig. 3e, g; n = 6), consistent with previous studies (6, 12). Importantly, administration of JAB0505 to ActA-mAb-treated hosts restored osteogenic differentiation of transplanted R206H-FAPs (Fig. 3e; n = 16), indicating that JAB0505 acts directly on R206H-FAPs and can replace the essential function of activin A in driving their skeletogenic differentiation. Notably, however, JAB0505 did not increase the quantity of bone generated by transplanted R206H-FAPs, regardless of whether ActA-mAb was co-administered to SCID hosts (Fig. 3e; n = 16). Production of peak heterotopic bone was modestly delayed when hosts were treated with JAB0505 (Supplemental Fig. 5; Day 14 vs. Day 10), a result that likely reflects both the receptor-blocking and weak receptor agonist activity of JAB0505, as in FOP mice.

In FOP mice, potentially responsive skeletal progenitors are scattered throughout and between muscles of the hindlimb. In contrast, progenitors of SCID hosts rarely contribute to HO because *Acvr1^R206H^* functions largely cell-autonomously (6), and JAB0505’s agonist activity is restricted to ACVR1(R206H)-expressing cells. That JAB0505 did not increase HO when a defined number of responsive cells was placed in a “non-recruitable” environment provides support for the relevance of enhanced and sustained progenitor cell recruitment as a mechanism of JAB0505 action following muscle injury of FOP mice. *A priori*, a direct positive effector function of JAB0505 on the proliferation of R206H-FAPs or their derivatives is a non-mutually-exclusive alternative model that could explain the exacerbating effects of JAB0505 on FOP mice. However, the inability of JAB0505 to enhance HO in this transplantation model argues against a primary role of JAB0505 in stimulating or sustaining proliferation of skeletal progenitors.

### Effects of JAB0505 on R206H-FAP population growth after muscle injury

To determine whether JAB0505 treatment causes alterations in R206H-FAP population growth dynamics in FOP mice, longitudinal live-animal luminescence imaging and flow cytometry were performed following muscle pinch injury. The Tie2-Cre driver was used to recombine the *Acvr1^tnR206H^* allele and the Cre-dependent luciferase reporter, *R26^luc^* (24). As Tie2-Cre is also expressed in hematopoietic cells and endothelium, these initial experiments provided a combined luminescent read-out of three critical cellular events following injury: immune cell infiltration, endothelial cell proliferation, and FAP population growth. After days 5 or 6, R206H-FAP-derived cartilage and bone also contribute to the luminescent signal, given the permanence of the Cre/loxP labeling system. The luminescent signal increased during the first few days following injury, and treatment with JAB0505 did not have a marked effect on apparent numbers of recombined cells at these early stages (Fig. 4a, b). However, whereas the apparent number of recombined cells in untreated FOP mice increased until day 5 and remained relatively constant thereafter, the luminescent signal from JAB0505-treated FOP mice continued to increase through 14-days post-injury (Fig. 4a, b). Quantification was not undertaken at day 21 because of the dampening of luminescence output by mineralized bone, particularly in JAB0505-treated mice. Interestingly, the luminescent signal in JAB0505-treated mice extended throughout most of the injured muscle, even in the first few days after injury (Fig. 4a), supporting the conclusion from histological analyses that a consequence of JAB0505 treatment is to engage muscle tissue over a much broader anatomical domain than in untreated FOP mice.

**Figure 4.**
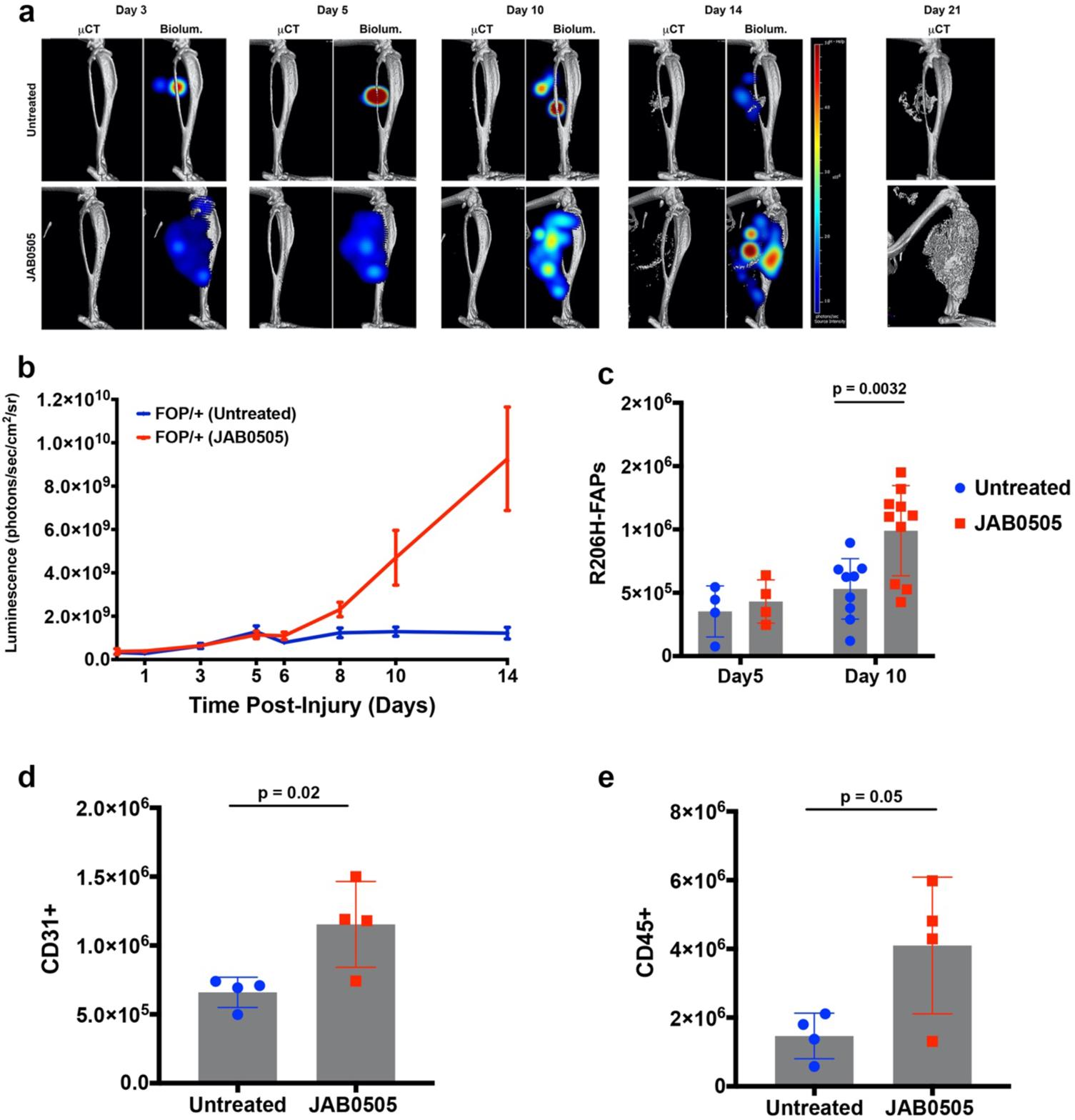
JAB0505 promotes sustained recruitment of R206H-FAPs to injury-induced lesions. (**a**) 3D tomographic bioluminescent source reconstruction following pinch injury of *Acvr1^tnR206H/+^*;*R26^Luc/+^*;Tie2-Cre FOP mice, with and without administration of JAB0505. Paired images show µCT alone (left panel) and µCT combined with the corresponding 3D bioluminescent reconstruction (right panel). For each experimental group, the same mouse is shown from days 3–21. Bioluminescent reconstruction was not performed at day 21 due to the dampening effect of dense bone on luminescent output. (**b**) Graphical representation of bioluminescent population dynamics of Tie2+ cells from *Acvr1^tnR206H/+^*;*R26^Luc/+^*;Tie2-Cre mice following pinch injury. Untreated, n = 16; JAB0505, n = 10. Error bars represent ± SEM. **(c)** Flow cytometry analysis to determine R206H-FAP cell number in injured *Acvr1^tnR206H/+^*;*R26^NG/+^*;Tie2-Cre which were either untreated (Day 5, n = 4; Day 10, n = 9) or administered JAB0505 at 10 mg/kg (Day 5, n = 4; Day 10, n = 10). p = 0.0032, two-way ANOVA. **(d, e)** Flow cytometry analysis to determine **(d)** CD31 cell number and **(e)** CD45+ cell number 10 days post-injury of *Acvr1^tnR206H/+^*;*R26^NG/+^*;Tie2-Cre mice that were either untreated (CD31+, n =4; CD45+, n=4) or administered JAB0505 at 10 mg/kg (CD31+, n =4; CD45+, n=4). Error bars represent t ± SD. Significance of **(d)** p = 0.02 for CD31+ cells and **(e)** p = 0.05 for CD34+ was determined by unpaired t-test.

As R206H-FAP-derived cartilage and bone contribute to the luminescent signal after the onset of skeletogenic differentiation, and because Tie2-Cre does not exclusively label FAPs, flow cytometry was employed to directly measure R206H-FAP numbers at the approximate onset of cartilage differentiation (day 5) and at an advanced stage of skeletogenic differentiation (day 10). R206H-FAPs were directly isolated from pinch-injured hindlimb muscles of *Acvr1^tnR206H/+^*;Tie2-Cre;*R26^NG/+^* mice as CD45-CD31-SCA1+GFP+tdTomato-cells (Supplemental Fig. 6; See Materials and Methods), where *R26^NG^* is a Cre-dependent GFP reporter (25) and tdTomato negativity identifies cells recombined at the *Acvr1^tnR206H^* locus (6). R206H-FAP numbers were similar between untreated and JAB0505-treated FOP mice at 5 days post-injury (Fig. 4c, n = 4). At day 10, the average number of R206H-FAPs was almost 2-fold greater in JAB0505-treated mice (Fig. 4c; p = 0.0032; n=10). Collectively, these data are consistent with the notion that JAB0505 exacerbates HO, in part, by acting to increase the number of R206H-FAPs that ultimately undergo skeletogenic differentiation. Although the increase in FAPs was not proportional to the profound increase in HO at endpoint, this may be explained by a sustained dynamic equilibrium between R206H-FAP population growth (by recruitment and proliferation, and possibly reduced apoptosis) and loss of these skeletal progenitors as they underwent commitment and differentiation to cartilage and bone. Finally, flow cytometric analysis at 10 days post-injury revealed a significantly higher total number of both CD31+ endothelial cells (Fig. 4d; n = 4, p = 0.02) and CD45+ hematopoietic cells (Fig. 4e; n = 4, p = 0.05) in injured muscle of JAB0505-treated mice. Whether the increase in CD45+ cells reflects a sustained inflammatory response caused directly or indirectly by JAB0505 treatment, and the extent to which this prolonged response drives the cellular changes correlated with worsened pathology, will require further investigation.

### Concluding remarks

The present findings show that JAB0505 inhibits ligand-dependent BMP signaling *in vitro* through wild-type ACVR1 and ACVR1(R206H) with reporter cell lines, but also functions as a weak agonist to activate signaling through ACVR1(R206H), ultimately driving growth and differentiation of pathogenic R206H-FAPs *ex vivo* and *in vivo*. The current data support the view that enhanced recruitment of R206H-FAPs, together with alterations in the balance between progenitor cell growth and differentiation, significantly extend the temporal and spatial domains of heterotopic bone formation following injury, ultimately leading to a dramatic increase in HO. Recent independent studies confirmed the ability of anti-ACVR1 antibodies to profoundly worsen HO in FOP mice, and further demonstrated that the HO-exacerbating effects were dependent on antibody bivalency and their ability to cluster ACVR1-containing receptor complexes (14). Understanding the pathological effects of anti-ACVR1 antibody action will require follow-up investigations that explore the connection between receptor clustering, downstream biochemical sequelae, and the tissue-level cellular effects reported here. Additional areas for further exploration include possible direct or indirect interactions of anti-ACVR1 antibodies with the immune system, which could increase or sustain inflammation, a known trigger for HO (19, 20). For example, interactions with immune cells through the Fc region of JAB0505 or other anti-ACVR1 antibodies might have immune-stimulatory effects that increase R206H-FAP recruitment to lesion formation. Indeed, the dysregulation of FAP population growth in FOP, which is exaggerated by JAB0505, may reflect disruption of immune system-imposed regulatory mechanisms that normally modulate FAP numbers after injury (26). Regardless of mechanism, it is clear that highly specific antibodies to ACVR1 have a profound ability to exacerbate disease progression in FOP mice. These findings raise serious safety and efficacy concerns for the use of anti-ACVR1 antibodies as a treatment modality to block ligand-receptor interactions and highlight the importance of using genetically and physiologically relevant mouse models and cell types to examine potential therapeutic candidates for FOP.

## Materials and Methods

### Reagents

Activin A, BMP6, BMP7, BMP9, and BMP10 were obtained from R & D Systems (Minneapolis, MN). The monoclonal antibody against activin A (ActA-mAb) was provided by Acceleron Pharma (Cambridge, MA), and was described previously (6).

### Discovery and optimization of the anti-ACVR1 monoclonal antibody

The Anti-ACVR1 monoclonal antibody was obtained from a phage display library of Fab fragments cloned from mice immunized with DNA encoding the ACVR1 extracellular domain (ECD). CD-1 and NZBWF/J mice were immunized by subcutaneous injection of ACVR1 ECD plasmid DNA suspended in sterile TE buffer supplemented with aSMAR DNA immunization adjuvant reagent (Antibody Research, St. Charles, MO) or with ACVR1 ECD plasmid DNA that was precipated onto gold beads and administered using a HELIOS Gene Gun (Bio-Rad).

Anti-ACVR1 specific sera titers from immunized mice were determined by ELISA. ELISA plates were coated with a mixture of recombinant human ACVR1 antigens consisting of ACVR1 ECD fused to human G1 Fc (Sino Biologicals,10227-H03B), human ACVR1 ECD fused to human G1 Fc (R&D Systems, 637-AR) and ACVR1 ECD (Creative Biomart, ACVR1-01H). Antibodies bound to ACVR1 were detected using goat anti-mouse IgG (H+L) – HRP (ThermoFisher Scientific, 62-6520).

A Fab phage display library was constructed from ACVR1 immunized, sera positive mice. Total RNA was isolated from lymph nodes and spleens. This was used as the template for cDNA synthesis using Superscript III First-Strand Synthesis System for RT-PCR (Life Technologies). The cDNA was used as the template for the amplification of heavy chain IgG1, IgG2a and light chain Ig kappa sequences. The antibody sequences were cloned in bulk into a phage display vector. The phagemid was used to transform XL1-blue E.coli cells to generate the anti-ACVR1 library.

The library was panned for two rounds on biotinylated ACVR1 ECD (Creative Biomart) that was conjugated to magnetic streptavidin beads (Dynabeads MyOne, 65601,ThermoFisher). Streptavidin beads without ACVR1 were used in deselection steps to remove non-specific binders. Phage eluted from ACVR1-coated beads were tested for binding to the ACVR1 ECD using an electrochemiluminescence immunoassay (Meso Scale Discovery, Rockville, Maryland). The variable region gene sequences of phage that bound to ACVR1 were cloned into mammalian expression plasmids for the expression of full length of mouse IgG1 kappa monoclonal antibodies.

Full-length antibodies were expressed in HEK293 cells and purified by protein A chromatography. Antibodies were tested for binding to the ACVR1 ECD by biolayer interferometry using an Octet HTX instrument (Pall/Forte Bio). Confirmed binders were evaluated for the ability to block BMP-mediated ACVR1 signaling in a cell-based reporter assay described below.

Affinity maturation of functional anti-ACVR1 antibodies was performed by mutating the complementary determining regions (CDRs) by saturation mutagenesis of each position of heavy chain CDRs 2 and 3 and light chain CDR3 independently using primers that randomly encode NNK at each specified codon where N=A/C/G/T and K=G/T. Mutated antibody sequences were pooled and cloned into a phagemid vector to generate an affinity maturation library. This phage library was panned for ACVR1 binders for four rounds of increasingly stringent binding conditions. Phage binders were sequenced, and convergent antibody sequences were cloned and expressed as murine IgGl mAbs. Antibodies were expressed and screened for binding to ACVR1 by BIAcore and evaluated for inhibition of BMP-mediated ACVR1 signaling using the BMP Response Element (BRE) luciferase reporter assays described below. The specificity of affinity matured antibodies for ACVR1 was confirmed by flow cytometry using ALK3 KO C2C12 cells.

The amino acid sequence of antibody JAB0505 has been published in U.S. Patent Application, “Anti-ALK2 Antibodies and Uses Thereof,” patent number PCT/US 20 19/064613, WO 2020/118011 A1.

### Wild-type C2C12 BMP-reporter cell line development

A BMP reporter plasmid was constructed by synthesizing a BMP-response element (BRE) described in (15). The BRE response element was inserted upstream of the minimal promotor region of plasmid pGL4.26 luc2/minP/Hygro (Promega E8441) that also encodes the luciferase reporter gene luc2. The sequence of the BRE element was confirmed by DNA sequencing. The plasmid was designated pGL4.26 BRE2. Vector construction and sequence confirmation were performed by GeneArt (Thermo Fisher Scientific).

Mouse C2C12 cells (ATCC CRL-1772) were transfected with plasmid pGL4.26 BRE2 using Lipofectamine 3000 (Thermo Fisher Scientific, L300015) according to manufacturer’s instructions. Transfected cells were cultured in 96-well plates in DMEM with high glucose and L-glutamine (ATCC 30-2002), 10% FBS (Tissue Culture Biology 101), and 200 µg/mL hygromycin (Thermo Fisher Scientific 10687-010). Stable clones were scaled up and cryopreserved.

Transfected C2C12 clones were evaluated for BMP-dependent luciferase expression by plating cells at 2.0 X 10^4^ viable cells per well in 96 well plates in selective media containing 1% FBS. Cells were stimulated with titrations of mouse BMP9 (R&D Systems 5566-BP-010) ranging from 10 ng/mL to 156 pg/mL. Luciferase expression was analyzed using the Dual-Glo Luciferase assay system (Promega, E2940). Luminescence was read with either a Spectromax Paradigm or Flexstation 3 plate reader. Clone 21E12 was found to produce the highest luminescence in response to BMP9. It was also found to stably respond to BMP9 after 41 generations and was selected for use in cell-based assays measuring inhibition of wild-type ACVR1.

### Development of ACVR1(R206H)-transfected C2C12-BRE-Luc reporter cell line

A synthetic DNA construct encoding mouse ACVR1 with the R206H mutation (G to A at nucleotide position 617) was assembled from synthetic oligonucleotides and PCR products and cloned into the pCMV6 entry mammalian expression vector (Origene), which also encodes a neomycin resistance marker. This vector was designated pCMV6-ALK2R206H. Vector construction and sequence verification were performed by GeneArt (Thermo Fisher Scientific).

C2C12 cells were transfected with plasmid pCMV6-ALK2R206H using Lipofectamine 3000 (Invitrogen #L300015) according to manufacturer’s instructions. Transfected cells were cultured in 96 well plates in selective media containing 400 µg/mL geneticin (Life Technologies #10131-027). Stable clones were scaled up in selective media and cryopreserved.

Clones were analyzed for increased cell surface expression of ACVR1 relative to wild-type C2C12 cells by flow cytometry. Cells were washed once in sterile PBS and stained with Zenon Alexa Fluor 647 (Molecular Probes #Z25608)-labeled polyclonal anti-human activin A receptor type 1 antibody (LSBio #LS-C122594). Labeled cells were analyzed by flow cytometry using a LSRII flow cytometer (BD Biosciences). Transfected clones that had a higher ACVR1 mean fluorescence intensity (MFI) than non-transfected C2C12 cells were transiently transfected with the BRE reporter plasmid pGL4.26 BRE2 described above using Lipofectamine LTX (Life Technologies #15338-100) according to the manufacturer’s instructions. Transfected cells were cultured overnight in media containing 1% FBS in 96 well plates. The following day, clones were treated with a dilution series of recombinant mouse BMP9 (R&D Systems #5566-BP-010) ranging from100 ng/ml to 0.1 ng/ml (4.1 nM to 4.1 pM) for 4 hours. Expression of luciferase was then determined using the Dual-Glo Luciferase Assay System (Promega #E2940). Luminescence was measured with a Spectramax Paradigm plate reader (Molecular Devices).

ACVR1(R206H)-transfected clones expressing luciferase constitutively, and at higher levels than wild-type C2C12 cells when stimulated with BMP9, were stably transfected with the BRE reporter plasmid using Lipofectamine 3000 (Invitrogen #L300015) according to manufacturer’s instructions. Transfected cells were cultured in 96-well plates in media containing 200 µg/mL hygromycin and 400 µg/mL geneticin and cryopreserved.

Clones resistant to both hygromycin and geneticin were stimulated with mouse BMP9 (R&D Systems #5566-BP-101) or mouse BMP4 (R&D Systems #5020-BP-010) in media containing 1% FBS and tested for expression of luciferase using the ONE-Glo™ + Tox Luciferase Reporter and Cell Viability Assay (Promega #E7120). Luciferase expression was measured using a Flexstation 3 plate reader (Molecular Devices). Clone 84C7 was found to have elevated basal constitutive BRE-luciferase expression and enhanced BMP9 and BMP4 BRE-luciferase expression compared to the non-ACVR1(R206H) transfected C2C12 BRE-luciferase clone 21E12. Clone 84C7 was selected for use in cell-based assays measuring inhibition of ACVR1(R206H).

### Generation of *Acvr1* and *Bmpr1a* (*Alk3*) knockouts in C2C12 BRE-reporter lines

The C2C12 BMP-response element (BRE)-reporter line 21E12 was used to generate *Acvr1* (*Alk2*)- and *Bmpr1a* (*Alk3*)-knockout (KO) BRE reporter lines. To generate *Acvr1* and *Bmpr1a* knockout clones, 21E12 cells were co-transfected with three plasmids; one encoding the Cas9 enzyme, one encoding a single guide RNA (sgRNA) specific for either the ACVR1 gene (targeting Exon 6 sgRNA 5’ -TGTAAGACCCCGCCGTCACCTGG -3’) or the *Bmpr1a* gene (targeting Exon 5 sgRNA 5’ -CATTATAGAAGAAGATGATCAGG -3’), and a reporter plasmid encoding GFP and H2KK sequences that require *Acvr1* sgRNA or *Bmpr1a* sgRNA-mediated frame shift for expression of GFP and H2KK, respectively. All plasmids were obtained from PNA Bio Inc. (Thousand Oaks, CA). Cells were transfected using Amaxa 4D-Nucleofector with X-unit (Lonza, Basel, Switzerland). Two days after transfection, cells were enriched magnetically by their expression of H2Kk using a MACSelect Transfected Cell Selection kit (Miltenyi Biotec, Sunnyvale, CA). Cas9-mediated indel formation at the *Acvr1* or *Bmpr1a* locus was confirmed by T7E1 mismatch analysis on the enriched cell populations (T7E1 Nuclease – New England Biolabs, #M0302S). H2Kk+ cells were cloned by limiting dilution in 96-well plates. Clones were screened for ACVR1 or ALK3 protein expression by flow cytometry with an anti-ACVR1 antibody (R&D Systems #AF637) and an anti-ALK3 antibody (Sino Biological #50078), respectively. *Acvr1* and ALK3 KO clones were confirmed by genomic sequencing and by loss of response to rhBMP9 (R&D Systems #3209-BP/CF) or rhBMP4 (R&D Systems #314-BP/CF), respectively.

### Characterization of BMP receptor expression in C2C12 cells

BMP receptor mRNA expression was profiled in C2C12 cells by RT-qPCR. C2C12 cells were cultured in DMEM with high glucose and L-glutamine (ATCC #30-2002),10% FBS (Tissue Culture Biology 101). Total RNA was isolated from C2C12 cells using a RNeasy Plus Universal kit (Qiagen #73404) and cDNA was synthesized using an iScript cDNA synthesis kit (BioRad #170-8890). The expression of *Acvrl1* (*Alk1*), *Acvr1* (*Alk2*), *Bmpr1a* (*Alk3*), *Bmpr1b* (*Alk6*), *Gapdh*, and β-actin was quantified by RT-qPCR using Taqman primer and probes (Thermo Fisher Scientific). Expression levels of BMP receptors were reported relative to GAPDH.

The cell surface expression of ACVR1 and ALK3 proteins on C2C12 cells was evaluated by flow cytometry. Cells were removed from adherent cultures using AssayComplete cell dissociation reagent (Eurofins #92-0009) and incubated with a polyclonal anti-ALK2 antibody (R&D Systems #AF637) or a polyclonal anti-ALK3 antibody (Sino Biological #50078-RP02). The isotype control cells were ChromPure goat IgG (Jackson ImmunoResearch #005-000-003) for anti-ALK2 and ChromPure rabbit IgG (Jackson ImmunoResearch #011-000-003) for anti-ALK3. Cells were washed and incubated with the secondary antibodies donkey anti-goat IgG, Alexa Fluor 488 (Thermo Fisher Scientific #A-11055) and goat anti-rabbit IgG Alexa Fluor 488 (Thermo Fisher Scientific #A-11070) for anti-ALK2 and anti-ALK3, respectively. Cells were washed again and analyzed with a BD LSR II flow cytometer (BD Biosciences).

### FOP mouse models and crosses to generate experimental mice

#### *Acvr1^tnR206H^* FOP mouse model

Generation of *Acvr1^tnR206H^* mice was previously described (6), and standard breeding schemes were used to produce mice that carry the Tie2-Cre driver (17) (B6.Cg-Tg(Tek-cre)1Ywa/J; JAX #008863) and the reporters, *R26^NG^* (25) (derived from *R26^NZG^*; FVB.Cg-*Gt(ROSA)26Sor^tm1(CAG-lacZ,-EGFP)Glh^*/J; JAX #012429) or *R26^luc^* (24) (FVB.129S6(B6)-*Gt(ROSA)26Sor^tm1(Luc)Kael^*/J; JAX #005125). Adult mice between 8- and 16-weeks-of-age were used for all experiments. Only female mice were used in flow cytometry analysis of cell populations to account for possible sex differences in cell population numbers. Male and female mice were used interchangeably in all other studies. Mice were genotyped by PCR and reporter fluorescence, as previously described (6, 12).

#### *Acvr1^FLEx(R206H)^* FOP mouse model and tamoxifen induction

The *Acvr1^FLEx(R206H)^* tamoxifen-inducible FOP mouse model was developed by genOway (Lyon, France). The FLEx system (27) is a Cre-dependent one-way genetic switch that allows for expression of a mutant form of a gene in place of its wild-type copy. Briefly, *Acvr1^FLEx(R206H)^* was designed such that mutant exon 5, which carries the 617G->A substitution and was inserted in reverse orientation in intron 5, is switched to sense orientation and replaces the wild-type exon by the action of Cre. The mutant gene is expressed under the control of the endogenous *Acvr1* promoter. Electroporation of the targeting vector into 129Sv embryonic stem (ES) cells, G418 selection, and validation of targeted clones followed routine methods. Prior to production of chimeric mice by blastocyst injection, validated ES clones were transfected with a plasmid encoding Flp recombinase to remove the Frt-flanked neo cassette present in the targeting vector. Two highly chimeric males were mated with wild-type 129 females to establish *Acvr1^FLEx(R206H)^* mouse lines. Mice were maintained as heterozygotes on a 129 background. Details of the targeting vector design and construction, and ES cell and mouse line validation methods are available upon request.

To produce experimental mice, heterozygous *Acvr1^FLEx(R206H)^* female mice were crossed with heterozygous B6.Cg-Tg(CAG-cre/Esr1*)5Amc/J (JAX #004682) males to generate mice that were heterozygous at both loci, referred to as *Acvr1^FLEx(R206H)/+^*;CAG-Cre^ERT2^. The Cre driver CAG-cre/Esr1 was selected to provide high levels of Cre-recombinase expression across most cell types, and its activity is tamoxifen dependent. Mice between 4- and 8-weeks-of-age were used, and males and females were randomly assigned to study groups.

To induce recombination at the *Acvr1^FLEx(R206H)^* locus, 75 mg/kg tamoxifen (Sigma-Aldrich, St. Louis, MO, U.S., #T5648) prepared as a 20 mg/mL stock in corn oil (Sigma-Aldrich, St. Louis, MO, U.S., #C8267) was administered to experimental mice for 5 consecutive days. A 5–7-day washout period was incorporated prior to muscle injury.

### Skeletal muscle injury, dosing regimen, and endpoints

#### *Acvr1*^FLEx(R206H)/+^_;_CAG-Cre^ERT2^ mice

FOP mice were randomized into treatment groups consisting of 7 mice per group or into the vehicle control group consisting of three animals. Injury was induced by a single intramuscular injection of 100 µL of 10 µM cardiotoxin (Sigma-Aldrich, St. Louis, MO, U.S., #C9759) into the gastrocnemius muscle while mice were under isoflurane anesthesia. Animals received a single 10 mg/kg dose of antibody JAB0505 by i.v. injection into the tail vein on day -1 relative to cardiotoxin injury. Heterotopic bone formation was visualized by µCT on days 7 (n=7), 14 (n=7), and 20 (n=4) post-cardiotoxin injury.

#### *Acvr1^tnR206H/+^*;Tie2-Cre mice

The gastrocnemius muscle of adult mice was injured by applying 3500 – 3700 grams of force with a Randall Selitto Paw Pressure Test Apparatus (IITC Life Science, Woodland Hills, CA). Care was taken to avoid incidental contact with the tibia and fibula, which was verified by µCT. Injuries were performed while mice were under isoflurane anesthesia. Treated mice received a single subcutaneous dose of ActA-mAb (10 mg/kg) and/or a single intraperitoneal dose of the anti-ACVR1 monoclonal antibody JAB0505 (10 mg/kg) on the day of injury. Histological analyses were performed on days 6 and 14 post-injury, flow cytometry analyses were performed on days 5 and 10 post-injury, and µCT imaging was conducted on days 14, 21, 28, and 35 post-injury.

To produce a sub-clinical injury that does not cause HO in this model, the tibialis anterior muscle was injected with 50 μL of 2.5% methylcellulose (Sigma) in sterile PBS. JAB0505 administration was as above. Methylcellulose-injected mice were imaged by µCT at 15 and 22 days after injury.

### µCT and HO quantification

µCT imaging of FLEx ACVR1(R206H);CAG-cre/Esr1 mice was performed using a Quantum FX µCT Cabinet X-Ray System (PerkinElmer, Waltham, MA, U.S.), with mice under isoflurane anesthesia and placed in a supine position for hindlimb imaging. Scan parameters included 90 kV voltage, an 80 µm voxel size and a 4.5-minute scan time. µCT imaging of Acvr1^tnR206H/+^;Tie2-Cre mice was performed using an IVIS SpectrumCT (Perkin-Elmer; Hopkinton, MA). All µCT images were taken using medium resolution acquisition mode (75 µm voxel size; estimated radiation dose of 132 mGy; 210 s scan time) with mice under isoflurane anesthesia. The µCT images were generated and HO volumes were quantified using 3D Slicer software (http://www.slicer.org), as previously described (6, 12). Imaging of SCID hosts following cell transplantation was performed similarly.

### *In vivo* bioluminescence imaging

Bioluminescence images were acquired using an IVIS SpectrumCT and analyzed with Living Image 4.5 software (Perkin-Elmer; Hopkinton, MA). *Acvr1^tnR206H/+^;R26^luc/+^*;Tie2-Cre mice were injected IP with D-luciferin (Perkin Elmer, Hopkinton, MA) at 150 mg/kg prior to bioluminescence imaging. The bioluminescent light emission plateau was empirically determined to be 14 –18 min after D-luciferin substrate injection. Animals were anesthetized using the built-in XGI-8 Gas Anesthesia System with oxygen containing 2% isoflurane and placed into the imaging chamber. To determine bioluminescent source depth and signal intensity at the source, µCT and 2D surface radiance bioluminescent imaging was followed by 3D Diffuse Luminescent Imaging Tomography (DLIT) reconstruction as previously described (12).

### Histology and immunohistochemistry

Tissues were fixed, decalcified, and processed for paraffin embedding as previously described (6, 12). Following de-paraffinization, immunohistochemistry for ACVR1 was performed using a Rabbit anti-ACVR1 (Abcam; #ab60157), as previously described (6). Hematoxylin, eosin, and Alcian blue staining was performed using standard methods.

### FAP isolation and culturing

Skeletal muscle tissue dissociation by enzymatic digestion and isolation of FAPs by fluorescence activated cell sorting (FACS) was conducted as previously described (6, 12, 18, 28) with the following modification; magnetic depletion of CD31+ and CD45+ cells prior to FACS was omitted, and CD31+ and CD45+ cells were detected and sorted with anti-CD31-bv711 (BD Biosciences; 1:800) and anti-CD45-bv711 (BD Biosciences; 1:500) antibodies. Sorting was performed on a FACS Aria II (BD Biosciences, Franklin Lakes, NJ) equipped with 407, 488, and 633 lasers.

FACS-isolated FAPs were grown on tissue culture flasks (Nunc, Rochester, NY) in Dulbecco’s Modified Eagle Medium (DMEM; Life Technologies, Carlsbad, CA) with 50 U/mL Penicillin and 50 μg/mL Streptomycin (Pen/Strep; Gibco, Billings, MT) and 20% Premium fetal bovine serum (FBS), Atlanta Biologicals; Lot #C19032) as previously described (6, 12). FAPs were maintained at 37°C in a humidified atmosphere at 5% CO_2_. Media was changed every other day. All experiments utilized FAPs that had undergone less than 7 population doublings (approximately 3 passages).

### Quantification of cell populations by flow cytometry

All muscle groups of the posterior, lower hindlimb were harvested from untreated and JAB0505-treated mice at days 5 and 10 after muscle injury, and prepared for flow cytometry as described above. For this study, FAPs were defined as CD31-CD45-SCA-1+ cells, the vast majority of which express the FAP marker, PDGFRα (6, 18, 29). FAPs that were recombined and unrecombined at the *Acvr1^tnR206H^* locus were identified by the absence or presence, respectively, of tdTomato-derived fluorescence. (6). Absolute cell numbers were quantified using Precision Count Beads (BioLegend). For quantification of CD31+ and CD45+ populations, cell suspensions were divided evenly and stained with cell-specific antibodies separately, as both antibodies were conjugated to the same fluorophore. Flow cytometry data was collected using the FACS Aria II (BD Biosciences, Franklin Lakes, NJ) and was analyzed using FlowJo software (v10.6.1).

### Osteogenic and chondrogenic assays

Chondrogenic and osteogenic assays were performed using FACS-isolated and expanded FAPs, as previously described (6, 12). In brief, for chondrogenic differentiation, FAPS were resuspended at 2 X 10^7^ cells/mL and plated in 10 µL high-density micromass dots onto 35 mm tissue culture dishes (Nunc) in DMEM/F12 media (Life Technologies) containing 5% FBS plus 1% Pen/Strep. After 10 days of culture, micromasses were fixed in 10% neutral-buffered formalin and stained with Alcian blue, to detect cartilage-specific proteoglycans, as previously described (30). For osteogenic differentiation, FAPs were plated at a starting density of 3 X 10^4^ cells/cm^2^ and grown for 8 days in DMEM (Life Technologies) containing 5% FBS plus 1% Pen/Strep. ALP activity was detected using a commercial kit according to the manufacturer’s recommendations (Sigma). For both chondrogenic and osteogenic assays, FAPs were treated continually with exogenous ligand or mAb at the indicated concentrations.

### Western blots

Near-confluent monolayer cultures of FACS-isolated FAPs in 6-well plates were serum starved for 2 hours and then incubated for 1 hour with or without ligands and antibodies at the indicated concentrations. In the JAB0505 + ActA-mAb condition, FAPs were incubated with ActA-mAb 10 minutes prior to, as well as during, the 1-hour incubation with JAB0505 to allow time to sequester endogenous activin A. Procedures for cell lysis, SDS-PAGE, electrophoretic transfer, antibody incubation, and washing were previously described (6). Total protein concentration was measured using the DC protein assay (Bio-Rad), and equal amounts of total protein were loaded per lane of the same gel. After the final wash, membranes were incubated in in SuperSignal™ West Pico PLUS Chemiluminescent Substrate (ThermoFisher) and imaged using the IVIS SpectrumCT (Perkin-Elmer). pSMAD1/5/8 and β-actin bands were imaged separately, as separate secondary antibody incubations of the same membrane were required.

### Cell transplantation and antibody treatment

1 X 10^6^ FACS-isolated and expanded FAPs were resuspended in 50 µL of ice-cold 1X Dulbecco’s Phosphate-Buffered Saline (DPBS; Gibco, Billings, MT) and injected into the pinch injured gastrocnemius muscle of SCID Hairless Outbred mice (SHO*-Prkdc^scid^Hr^hr^*; Charles River) as previously described (6, 12). Treated SCID host mice received a single subcutaneous dose of ActA-mAb (10 mg/kg) and/or a single intraperitoneal dose of the anti-ACVR1 monoclonal antibody, JAB0505 (10 mg/kg), on the day of injury.

### Statistical analysis

Statistical analysis was performed using GraphPad Prism (GraphPad, La Jolla, CA). All numerical values are presented as mean values ± the standard error of the mean or standard deviation. Unpaired t-test and one or two-way ANOVA was used to determine significance, as described in the corresponding figure legends. Differences were considered significant at p ≤ 0.05.

## Acknowledgements

We thank Acceleron Pharma for the anti-activin A antibody, Michael J. Schneider for technical assistance with cell culture differentiation assays, Dr. Wu He, Facility Director of the Flow Cytometry Facility, for advice and technical assistance with FACS, and Anjli Kukreja, Sylvia Vong, and Vineeth Murali for technical support in developing cell based assays and characterizing anti-ACVR1 antibodies. This work was funded by a grant from the NIH (R01AR072052) to DJG, and by a sponsored research agreement between Alexion Pharmaceuticals and the University of Connecticut.

## Literature Cited

1. Pignolo RJ et al. The Natural History of Flare-Ups in Fibrodysplasia Ossificans Progressiva (FOP): A Comprehensive Global Assessment. Journal of bone and mineral research : the official journal of the American Society for Bone and Mineral Research 2016;31(3):650–656.

2. Kaplan FS et al. Early mortality and cardiorespiratory failure in patients with fibrodysplasia ossificans progressiva. The Journal of bone and joint surgery. American volume 2010;92(3):686–691.

3. Shore EM et al. A recurrent mutation in the BMP type I receptor ACVR1 causes inherited and sporadic fibrodysplasia ossificans progressiva. Nature Genetics 2006;38(5):525–527.

4. Hatsell SJ et al. ACVR1R206H receptor mutation causes fibrodysplasia ossificans progressiva by imparting responsiveness to activin A. Science translational medicine 2015;7(303):303ra137–303ra137.

5. Hino K et al. Neofunction of ACVR1 in fibrodysplasia ossificans progressiva. [Internet]. Proceedings of the National Academy of Sciences of the United States of America 2015;112(50):15438–15443.

6. Lees-Shepard JB et al. Activin-dependent signaling in fibro/adipogenic progenitors causes fibrodysplasia ossificans progressiva. Nat Commun 2018;9(1):471.

7. Wang H, Shore EM, Pignolo RJ, Kaplan FS. Activin A amplifies dysregulated BMP signaling and induces chondro-osseous differentiation of primary connective tissue progenitor cells in patients with fibrodysplasia ossificans progressiva (FOP). Bone 2017;109:218–224.

8. Billings PC et al. Dysregulated BMP signaling and enhanced osteogenic differentiation of connective tissue progenitor cells from patients with fibrodysplasia ossificans progressiva (FOP). Journal of bone and mineral research : the official journal of the American Society for Bone and Mineral Research 2008;23(3):305–313.

9. Haupt J, Xu M, Shore EM. Variable signaling activity by FOP ACVR1 mutations. Bone 2018;109:232–240.

10. Culbert AL et al. Alk2 regulates early chondrogenic fate in fibrodysplasia ossificans progressiva heterotopic endochondral ossification. Stem Cells 2014;32(5):1289–1300.

11. Hildebrand L, Stange K, Deichsel A, Gossen M, Seemann P. The Fibrodysplasia Ossificans Progressiva (FOP) mutation p.R206H in ACVR1 confers an altered ligand response. Cell Signal 2017;29:23–30.

12. Lees-Shepard JB et al. Palovarotene reduces heterotopic ossification in juvenile FOP mice but exhibits pronounced skeletal toxicity. eLife 2018;7:305.

13. Dey D et al. Two tissue-resident progenitor lineages drive distinct phenotypes of heterotopic ossification [Internet]. Science translational medicine 2016;8(366):366ra163–366ra163.

14. Aykul S et al. ACVR1 antibodies exacerbate heterotopic ossification in fibrodysplasia ossificans progressiva (FOP) by activating FOP-mutant ACVR1. bioRxiv 2021; doi:10.1101/2021.07.18.452865

15. Yadav PS, Prashar P, Bandyopadhyay A. BRITER: A BMP Responsive Osteoblast Reporter Cell Line. Plos One 2012;7(5):e37134.

16. Sanchez-Duffhues G, Williams E, Goumans M-J, Heldin C-H, Dijke Pten. Bone morphogenetic protein receptors: Structure, function and targeting by selective small molecule kinase inhibitors. Bone 2020;138:115472.

17. Kisanuki YY et al. Tie2-Cre transgenic mice: a new model for endothelial cell-lineage analysis in vivo. Developmental Biology 2001;230(2):230–242.

18. Wosczyna MN, Biswas AA, Cogswell CA, Goldhamer DJ. Multipotent progenitors resident in the skeletal muscle interstitium exhibit robust BMP-dependent osteogenic activity and mediate heterotopic ossification. [Internet]. Journal of bone and mineral research : the official journal of the American Society for Bone and Mineral Research 2012;27(5):1004–1017.

19. Genêt F et al. Neurological heterotopic ossification following spinal cord injury is triggered by macrophage-mediated inflammation in muscle. J Pathol 2015;236(2):229–240.

20. Convente MR et al. Depletion of Mast Cells and Macrophages Impairs Heterotopic Ossification in an Acvr1 R206H Mouse Model of Fibrodysplasia Ossificans Progressiva. J Bone Miner Res 2018;33(2):269–282.

21. Rodgers JT et al. mTORC1 controls the adaptive transition of quiescent stem cells from G0 to GAlert. Nature 2014;510(7505):393–396.

22. Nickel J, Mueller TD. Specification of BMP Signaling. Cells 2019;8(12):1579.

23. Lounev VY et al. Identification of progenitor cells that contribute to heterotopic skeletogenesis. [Internet]. The Journal of bone and joint surgery. American volume 2009;91(3):652–663.

24. Safran M et al. Mouse reporter strain for noninvasive bioluminescent imaging of cells that have undergone Cre-mediated recombination. Mol Imaging 2003;2(4):297–302.

25. Yamamoto M et al. A multifunctional reporter mouse line for Cre- and FLP-dependent lineage analysis. Genesis (New York, N.Y. : 2000) 2009;47(2):107–114.

26. Lemos DR et al. Nilotinib reduces muscle fibrosis in chronic muscle injury by promoting TNF-mediated apoptosis of fibro/adipogenic progenitors. Nat Med 2015;21(7):786–794.

27. Schnütgen F et al. A directional strategy for monitoring Cre-mediated recombination at the cellular level in the mouse. Nat Biotechnol 2003;21(5):562–565.

28. Biswas AA, Goldhamer DJ. FACS Fractionation and Differentiation of Skeletal-Muscle Resident Multipotent Tie2+ Progenitors. Methods Mol Biol 2016;1460(Chapter 18):255–267.

29. Joe AWB et al. Muscle injury activates resident fibro/adipogenic progenitors that facilitate myogenesis. Nat Cell Biol 2010;12(2):153–163.

30. Gay SW, Kosher RA. Uniform cartilage differentiation in micromass cultures prepared from a relatively homogeneous population of chondrogenic progenitor cells of the chick limb bud: Effect of prostaglandins. J Exp Zool 1984;232(2):317–326.

