## Supplemental Figures 1-6 for "An anti-ACVR1 antibody exacerbates heterotopic ossification by fibro/adipogenic progenitors in fibrodysplasia ossificans progressiva mice"

### Supplemental Figure 1

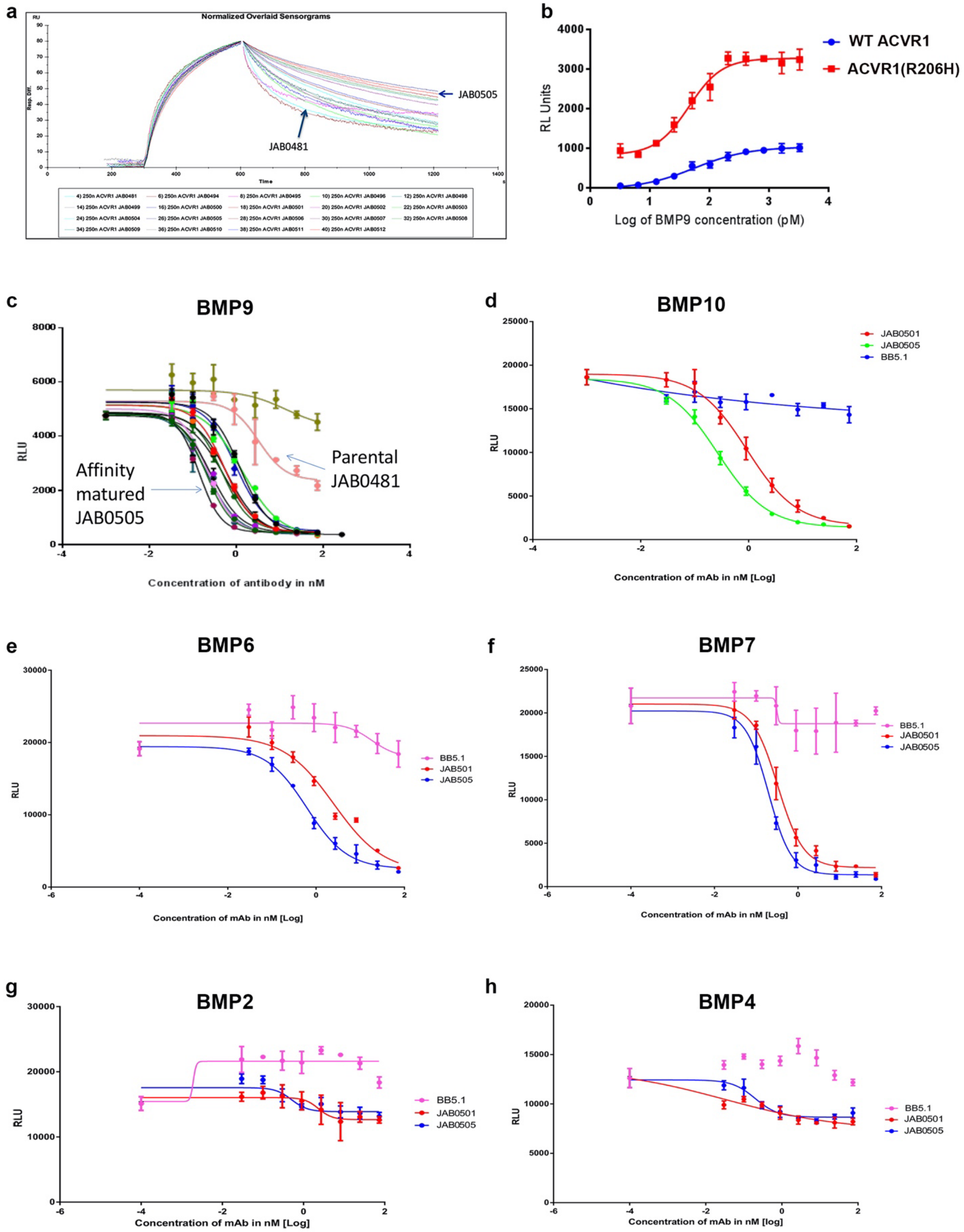

**Supplemental Figure 1. Characterization of JAB0505. (a)** Normalized sensorgram overlay displaying binding rate constants and equilibrium dissociation rate constants of affinity matured variants of the parental anti-ACVR1 antibody JAB0481. **(b)** Wild-type and ACVR1(R206H)-overexpressing C2C12-BRE-Luc cells were treated with BMP9 to establish a dose response. **(c)** C2C12-BRE-Luc cells were treated with BMP9 together with anti-ACVR1-antibodies. The IC50 of JAB0505 was 140 pM. **(d-h)** Anti-ACVR1 antibodies JAB0505 and JAB0501 effectively block BMP-signal activation by **(d)** BMP10, **(e)** BMP6, and **(f)** BMP7 in C2C12-BRE-Luc cells expressing wild-type ACVR1. Anti-ACVR1 antibodies JAB0505 and JAB0501 exhibit weak capacity to inhibit **(g)** BMP2 or **(h)** BMP4 in C2C12-BRE-Luc cells stably overexpressing ACVR1(R206H). BB5.1 is an IgG isotype control antibody.

**Supplemental Figure 2**

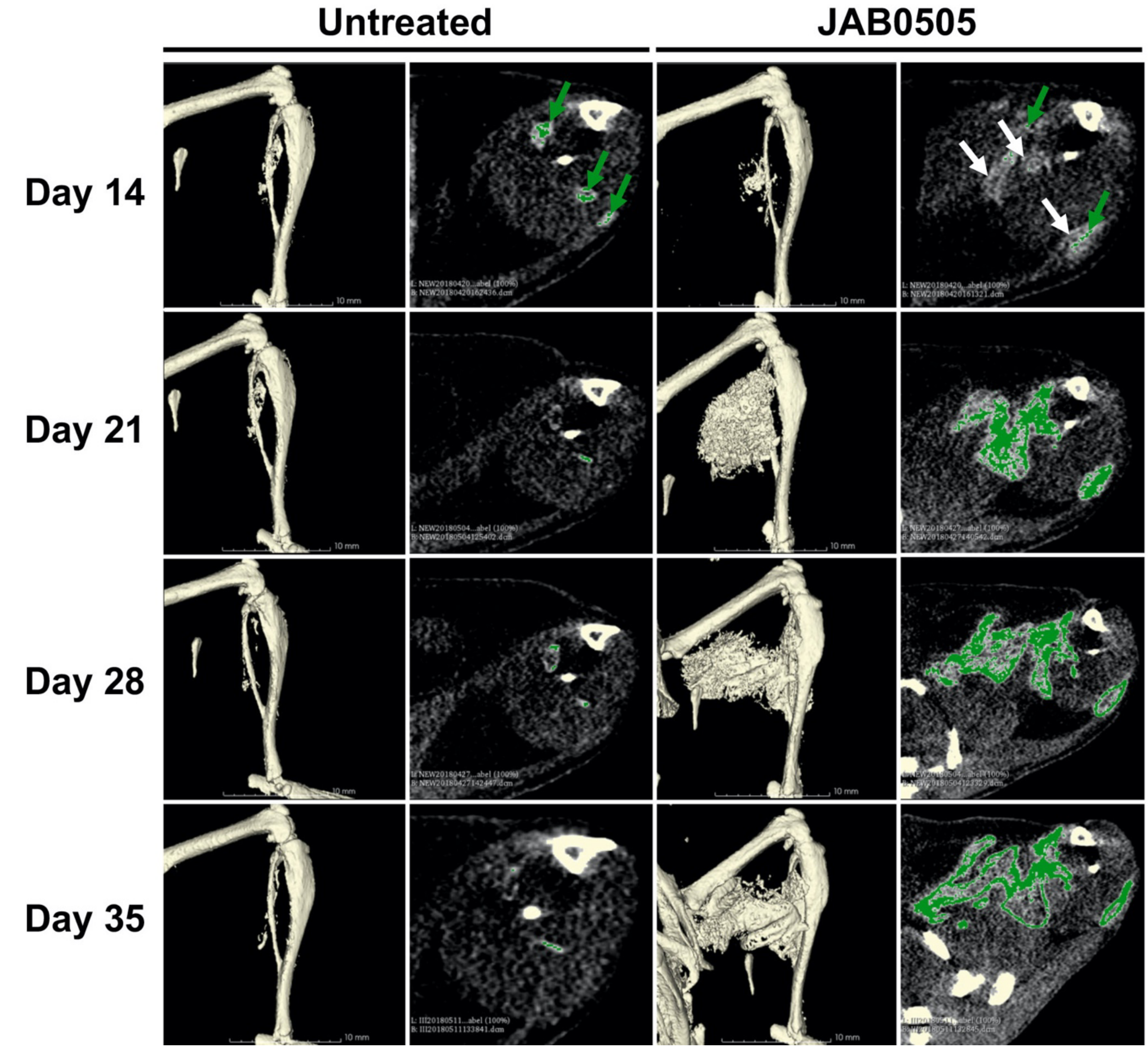


**Supplemental Figure 2. Progression of HO in *Acvr1*$^{tnR206H/+}$;Tie2-Cre FOP mice with and without JAB0505.** Paired single transverse slice and 3D reconstructed µCT images of the distal hindlimb of *Acvr1*$^{tnR206H/+}$;Tie2-Cre mice at the indicated times after hindlimb muscle pinch injury with and without administration of JAB0505. Radio-opaque lesional tissue at (HO: green arrows) or below (white arrows) the threshold set for quantification of mineralized bone are indicated. At day 14, JAB0505-treated mice commonly had extensive accumulations of tissue that was detectable, but below the threshold set for quantification of mineralized bone.

**Supplemental Figure 3**

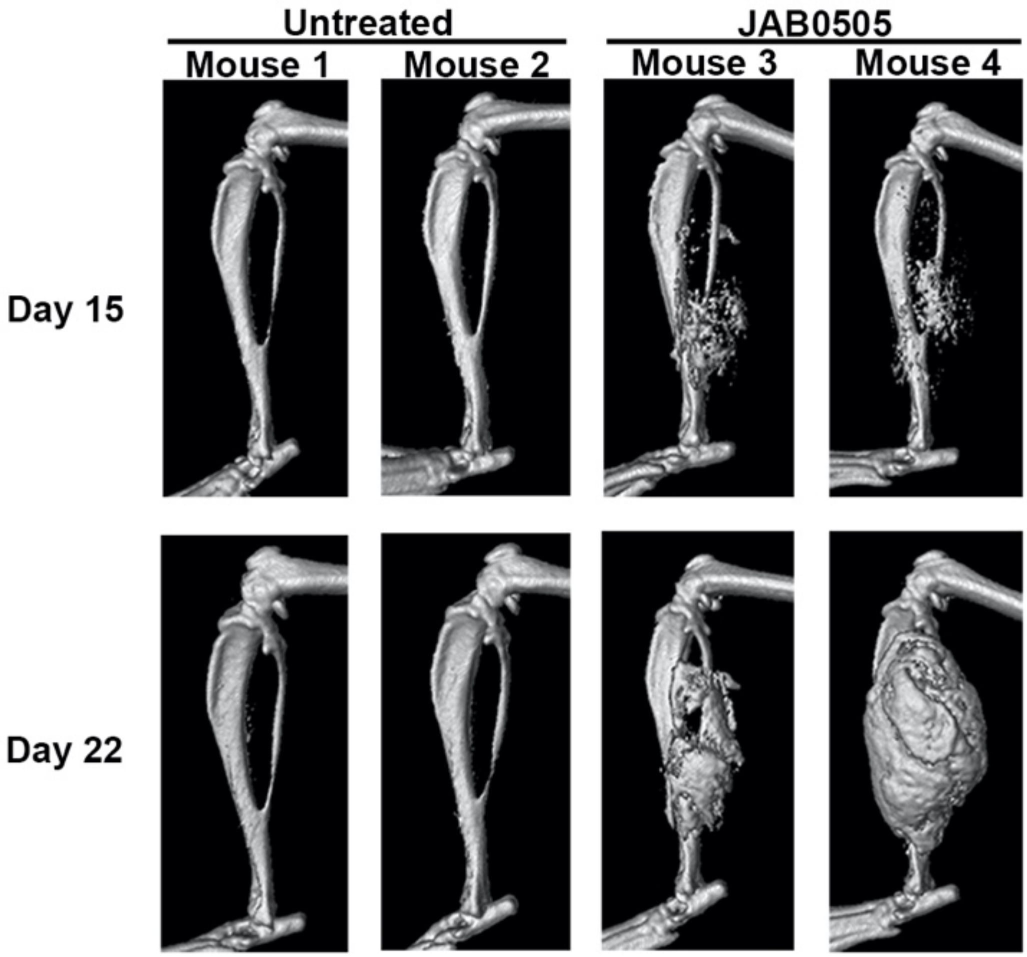


**Supplemental Figure 3. JAB0505 lowers the injury threshold necessary to cause HO in *Acvr1*$^{tnR206H/+}$;Tie2-Cre FOP mice.** μCT images of the distal hindlimbs of *Acvr1*$^{tnR206H/+}$;Tie2-Cre FOP mice at indicated time points after injection of 50 μL of 2.5% methylcellulose into the tibialis anterior muscle, with and without administration of 10 mg/kg JAB0505 (untreated, n=2; JAB0505-treated, n=2). A lateral view of the left hindlimb of each mouse is shown. Contralateral hindlimbs (not shown) received equivalent injuries and formed comparable HO.

**Supplemental Figure 4**

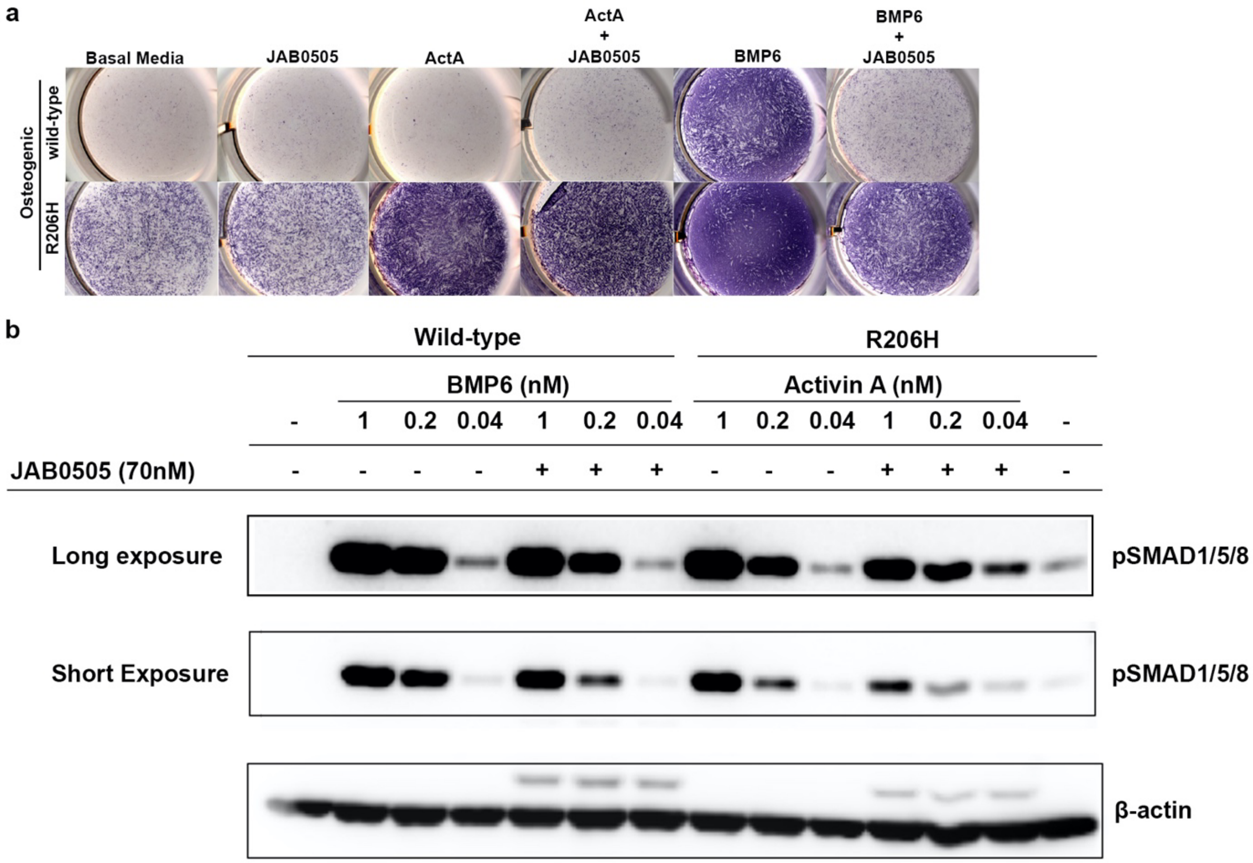


**Supplemental Figure 4. Effects of JAB0505 on osteogenic differentiation and ACVR1 signal activation. (a)** Osteogenic differentiation, as assessed by ALP staining (purple) and of wild-type FAPs and R206H-FAPs (R206H) cultured with or without 25 ng/mL (~1 nM) activin A or BMP6. JAB0505 was used at 10 μg/mL (~70 nM) **(b)** Western blot of SMAD1/5/8 phosphorylation (pSMAD1/5/8) in response to activin A or BMP6 with or without JAB0505. Shorter and longer exposures for pSmad1/5/8 are provided to better show the signal dynamic range and to more accurately visualize the degree to which JAB0505 reduces pSMAD1/5/8 levels (shorter exposure). β-actin was used as a loading control.

**Supplemental Figure 5**

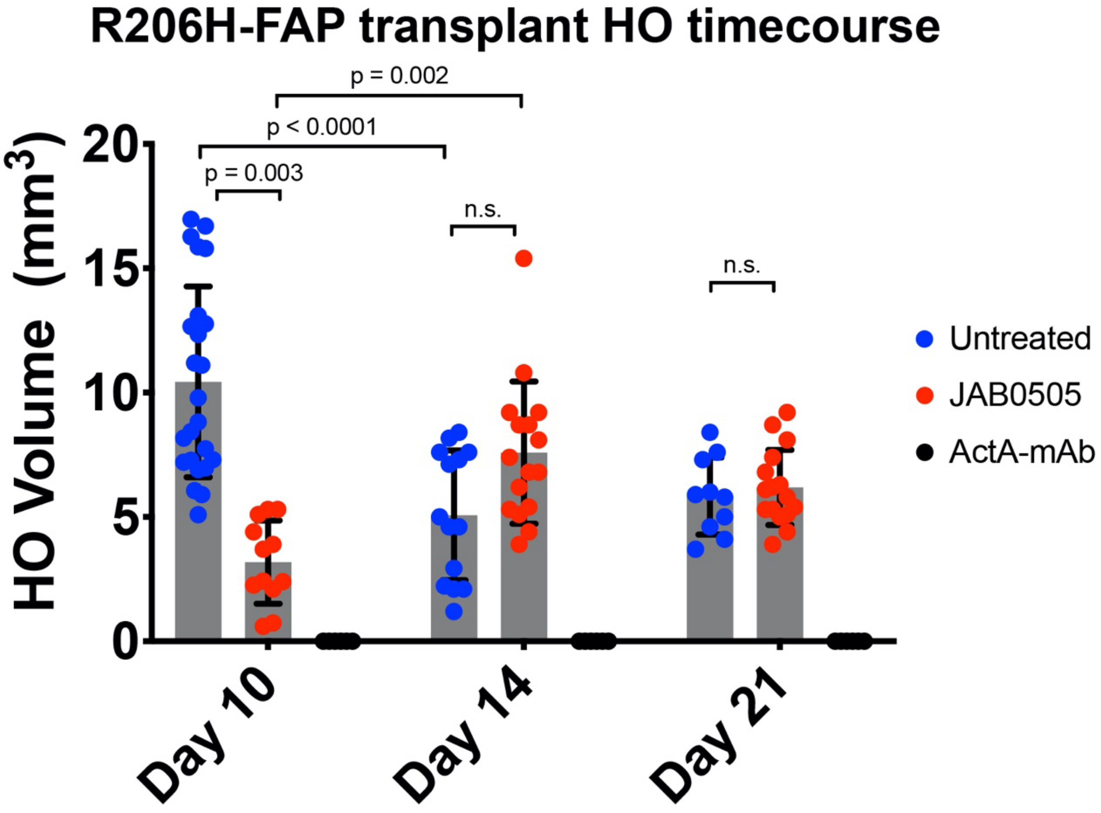


**Supplemental Figure 5. JAB0505 slows the time course of HO by transplanted R206H-FAPs.** Quantification of HO volume as measured by µCT of the distal hindlimb post-transplantation of R206H-FAPs into the pre-injured gastrocnemius of SCID hosts. Untreated: day 10, n = 24; day 14, n = 14; day 21, n = 10. JAB0505 (10 mg/kg): day 10, n = 12, day 14, n = 16, day 21, n = 16. ActA-mAb (10 mg/kg): n = 6 at all time points. Bars represent ± SD. Significance was determined by two-way ANOVA.

### Supplemental Figure 6

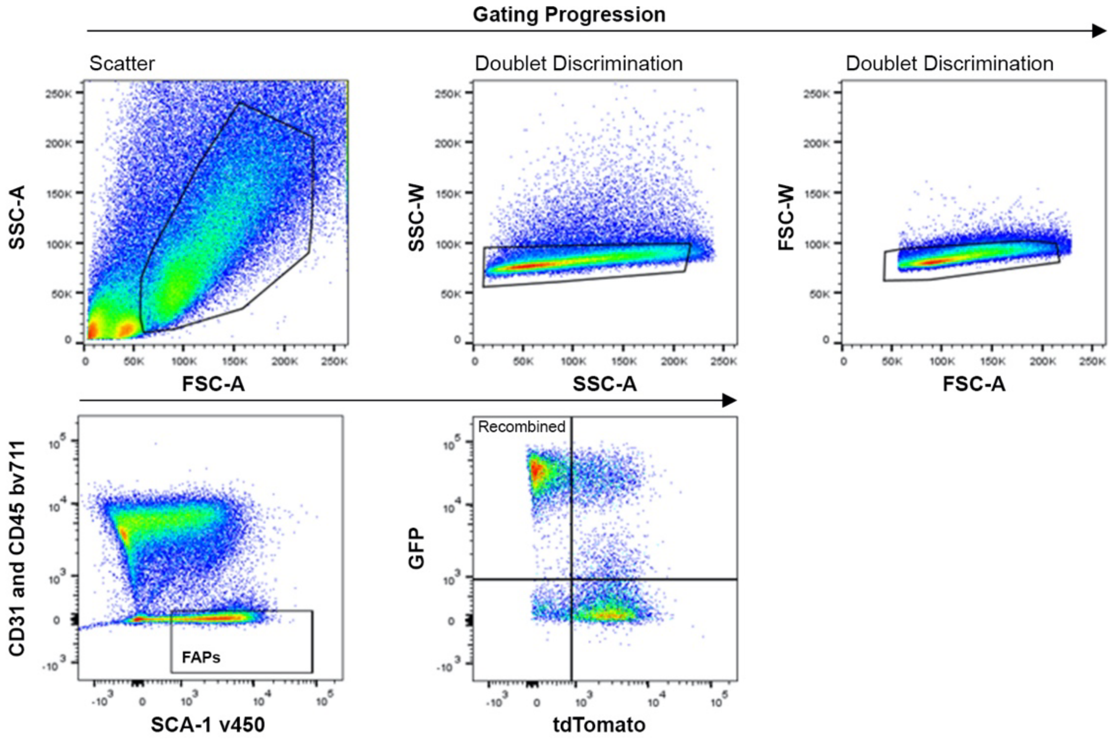


**Supplemental Figure 6. FACS gating strategy for isolation of R206H- FAPs and quantification of FAPs, CD31+, and CD45+ populations.** Plots shown are representative of the gating progression used for cell population quantification and sorting of FAPs for downstream assays. Total mononuclear cells from enzymatically digested, injured skeletal muscle were gated for side/forward scatter to eliminate debris and then for doublet discrimination. When sorting FAPs, bv711 was used to gate out both CD31+ and CD45+ populations. FAPs from control muscle were identified as the CD31-CD45-SCA-1+ fraction, while recombined FAPs from *Acvr1*$^{R206H/+}$;Tie2-Cre muscle were identified as the CD31-CD45-SCA-1+GFP+tdTomato- fraction. Equivalent sample subfractions were used for quantification of CD31+ and CD45+ populations (not shown), as bv711-congugated antibodies were used to label both populations.
